# Fine-grained habitat-associated genetic connectivity in an admixed population of mussels in the small isolated Kerguelen Islands

**DOI:** 10.1101/239244

**Authors:** Christelle Fraïsse, Anne Haguenauer, Karin Gérard, Alexandra Anh-Thu Weber, Nicolas Bierne, Anne Chenuil

## Abstract

Reticulated evolution -i.e. secondary introgression / admixture between sister taxa-is increasingly recognized as playing a key role in structuring infra-specific genetic variation and revealing cryptic genetic connectivity patterns. When admixture zones coincide with ecological transitions, the connectivity patterns often follow environmental variations better than distance and introgression clines may easily be confounded with local adaptation signatures. The Kerguelen mussels is an ideal system to investigate the potential role of admixture in enhancing micro-geographic structure, as they inhabit a small isolated island in the Southern Ocean characterized by a highly heterogeneous environment. Furthermore, genomic reticulation between Northern species (*M. edulis, M. galloprovincialis* and *M. trossulus*) and Southern species (*M. platensis*: South America and the Kerguelen Islands; and *M. planulatus*: Australasia) has been suspected. Here, we extended a previous analysis by using targeted-sequencing data (51,878 SNPs) across the three Northern species and the Kerguelen population. Spatial structure in the Kerguelen was then analyzed with a panel of 33 SNPs, including SNPs that were more differentiated than the genomic average between Northern species (i.e., ancestry-informative SNPs). We first showed that the Kerguelen lineage splitted very shortly after *M. edulis* and *M. galloprovincialis* initiated speciation, and it subsequently experienced admixture with the three Northern taxa. We then demonstrated that the Kerguelen mussels were significantly differentiated over small spatial distance, and that this local genetic structure was associated with environmental variations and mostly revealed by ancestry-informative markers. Simulations of admixture in the island highlight that genetic-environment associations can be better explained by introgression clines between heterogeneously differentiated genomes than by adaptation.

## Introduction

Marine species with planktonic larvae are high-dispersal organisms, with large effective population sizes and living in a highly connected environment (Cowen & Sponaugle 2009), therefore they generally show low levels of genetic differentiation across their species ranges (Palumbi 1992). Nevertheless, there is some evidence that micro-geographic genetic-environment associations occur at specific loci in marine species, such as barnacles (Schmidt & Rand 1999), mussels (Koehn *et al.* 1980) or Atlantic killifishes (Reid *et al.* 2017), despite genome-wide genetic homogeneity. These locus-specific associations are traditionally expected in regions of the species range where environmental gradients promote local adaptation of specific populations (Schmidt *et al.* 2008). Alternatively, genetic-environment associations can be observed where semi-permeable genetic backgrounds form a contact zone that coincides with an ecological boundary (Bierne *et al.* 2011). In this case, reproductive isolation genes become locally coupled with local adaptation genes and linked neutral markers, and the two genetic backgrounds become associated with different habitats. In addition, introgression creates a departure from equilibrium situations in which the influx of heterospecific alleles from one genetic background generates a gradient in allele frequencies within the other genetic background, revealing cryptic connectivity patterns (Gagnaire *et al.* 2015). Gradients of introgression may easily be confounded with local adaptation signatures, especially when environmental variables are better proxy of the connectivity matrix than geographic distance. There are two reasons for that: (i) the environment can have a direct effect on connectivity (e.g. landscape resistance model,Zeller *et al.* 2012), or (ii) the effect of a barrier to dispersal can be better described by one of the many scored environmental variables that can incidentally or truly coincide with it, than by distance. In the sea, regions of little water mixing are inevitably both ecotones and barrier to larval dispersal. Most importantly, the reduction in gene flow between genetic backgrounds is expected to be visible only on a subset of markers either because localized at an intermediate linkage map distance to reproductive isolation genes (Gagnaire *et al.* 2015) or because of heterogeneous differentiation produced by linked selection (Simon and Duranton 2018). This potential effect of introgression on our capacity to detect connectivity breaks in apparently well-mixed populations is of central concern to conservation and species management (Gagnaire *et al.* 2015).

*Mytilus* mussels are an excellent system to address these issues, because they are subdivided into partially reproductively isolated species. Moreover, a recent study based on FST genome scans and gene genealogies demonstrated that population-specific introgression is widespread, and it is the primary cause of outlying levels of genetic differentiation between conspecific populations (Fraïsse *et al.* 2016). *Mytilus* mussels have an antitropical distribution, i.e. they occur in high latitudes of the Northern and Southern Hemispheres, as a result of transequatorial migration during the Pleistocene (Hilbish *et al.* 2000; Gérard *et al.* 2008). In the North, *M. edulis* and *M. galloprovincialis* are closely-related species which started to diverge about 2.5 mya (Roux *et al.* 2014), while *M. trossulus* is an outgroup to them with a divergence dated at 3.5 mya (Rawson & Hilbish 1995). The three species have experienced a complex history of divergence punctuated by periods of gene flow (Roux *et al.* 2014; Fraïsse et al. 2018); and nowadays they display hybrid zones where their ranges overlap (Skibinski *et al.* 1983; Väinölä & Hvilsom 1991; Bierne *et al.* 2003). In the South, a re-evaluation of allozyme data and a review of the results obtained with mtDNA and two nuclear DNA markers (Borsa *et al.* 2012) encouraged to group Southern mussels in two different taxa, namely *M. platensis* for those related to *M. edulis* (the South American and Kerguelen mussels), and *M. planulatus* for those related to *M. galloprovincialis* (the Australasian mussels). The presence of a mitochondrial clade endemic to the Southern Ocean further suggests Southern mussels are native rather than introduced by human-mediated activities (Hilbish *et al.* 2000; Gaitán-Espitia *et al.* 2016).

In the Southern Indian ocean, the isolated Kerguelen Islands harbor *M. platensis* mussels which are polymorphic for allozyme alleles characteristic of all three Northern species (Blot *et al.* 1988), although they are most similar to *M. edulis* at a few allozyme loci (McDonald *et al.* 1991). Further analyses with nuclear markers strengthened the view of a mixed genome ancestry of the Kerguelen mussels (Borsa *et al.* 2007): at Glu-5′, a Northern diagnostic marker, mussels carry a heterospecific polymorphism (*M. edulis* / *M. galloprovincialis*). These preliminary results suggest either that reproductive isolation genes responsible of the interspecific barrier in the North were not yet evolved at the time of admixture (if any) in the Kerguelen, or that isolation is not as strong in the demographic, ecological and genetic context of the Kerguelen Islands as it is in the Northern Hemisphere hybrid zones.

The geomorphology of the Kerguelen Islands has been shaped by volcanic activity and glacial erosion which resulted in a carved coast with sheltered bays and fjords (Gérard *et al.* 2015). Recently, Gérard *et al.* (2015) have investigated the genetic-environment associations in the island with four nuclear markers (mac-1, Glu-5′, EFbis and EFprem’s) and a mitochondrial gene (COI). Only Glu-5′ revealed significant genetic differentiation among and within geographic regions, and between habitats, despite the very small geographical scale. This pattern is striking as intraspecific genetic variation rarely shows significant differentiation even over large distances in *Mytilus* (e.g. within the Mediterranean Sea, Fraïsse *et al.* 2016). In particular, allele frequencies at Glu-5′ were associated with the presence/absence of the kelp *Macrocystis* in the island, which serves as substrate and refuge for many species of molluscs, including mussels (Adami & Gordillo, 1999). As such, local adaptation was invoked to explain the fine-scale maintenance of polymorphism at Glu-5’, although alternative interpretations involving admixture could not have been refuted.

Our aim was to investigate whether reticulate evolution actually contributed to micro-geographic structure in the Kerguelen islands. To this end, we ask the following questions: (i) can we detect footprints of introgression from Northern species and disentangle them from incomplete lineage sorting (i.e. shared ancestral polymorphism)?; (ii) is there genetic differentiation at fine scale in the island?; (iii) is there association of genetic differentiation with environmental variables and/or geography?; (iv) is the signal observed in (ii) and (iii) genome-wide or concentrated on ancestry-informative loci and (v) does introgression contribute to this fine-scale genetic differentiation? Based on published genotyping-by-sequencing (GBS) data of the three Northern species (Fraïsse *et al.* 2016) and new GBS data of a sample from a single Kerguelen population, we found that the Kerguelen Islands harbor a Southern lineage of mussels, related to *M. edulis,* and that was subsequently admixed with Northern species. The recent divergence between the Northern species and the Southern lineage resulted in non-negligible incomplete lineage sorting of two types: (i) an ancient incomplete lineage sorting that resolved into multifarious topologies among the four taxa, and (ii) ongoing sorting of lineages (i.e. true shared ancestral polymorphism), which results in unresolved relationships. However, we also robustly detected past introgression events between Northern and Southern mussels by: (i) testing for admixture with genome-wide allele frequency data, (ii) reconstructing gene genealogies at a small chromosomal scale and (iii) inferring their divergence history from the joint site-frequency spectrum. Based on a new panel of SNPs genotyped on 695 mussels across 35 sites in the Kerguelen, we then confirmed using redundancy analysis (RDA) a significant fine-scale genetic differentiation between sites associated with environmental variables. Notably, we found that loci with strong genetic-environment association also tended to be among the most ancestry-informative markers in the *Mytilus spp*. While we cannot refute whether local adaptation has played a role here, simulations of admixture in the island highlights the importance of introgression for the current and local genetic structure of the Kerguelen mussels.

## Materials and Methods

### Genotyping-by-sequencing of the *Mytilus* spp

We used samples collected from nine localities in the Northern Hemisphere (Figure 1, Supp. Info. M&M and Table S1) to investigate the patterns of admixture between Northern and Southern genetic backgrounds in the Kerguelen Islands. The genetic composition of these samples has been analysed in Fraïsse *et al.* (2016) with target enrichment sequencing of bacterial artificial chromosomes (BAC) and cDNA sequences (see Supp. Info. M&M for details). The Northern samples have been shown to be representative of monospecific panmictic patches of the *Mytilus edulis* species complex, which comprises three species that hybridize at several places in the Northern Hemisphere: *M. galloprovincialis*, *M. edulis* and *M. trossulus*. In addition to these previously published samples, eight individuals from the Kerguelen Islands (Baie de la Mouche, Table S1) were included in the target enrichment experiment. These individuals were treated together with the Northern samples following the genotyping-by-sequencing (GBS) method described in Fraïsse *et al.* (2016)(see Supp. Info. M&M for details). The final dataset across the ten localities consisted of 1,269 reference sequences (378 BAC contigs that come from a pool of 224 unlinked clones, and 891 cDNA contigs that correspond to unlinked coding sequences of known-functions or randomly selected) and 129,346 SNPs. DNA sequences and VCF files including GBS genotypes are available on Dryad doi: 10.5061/dryad.6k740 (Fraïsse *et al.* 2016).

**Figure 1.**
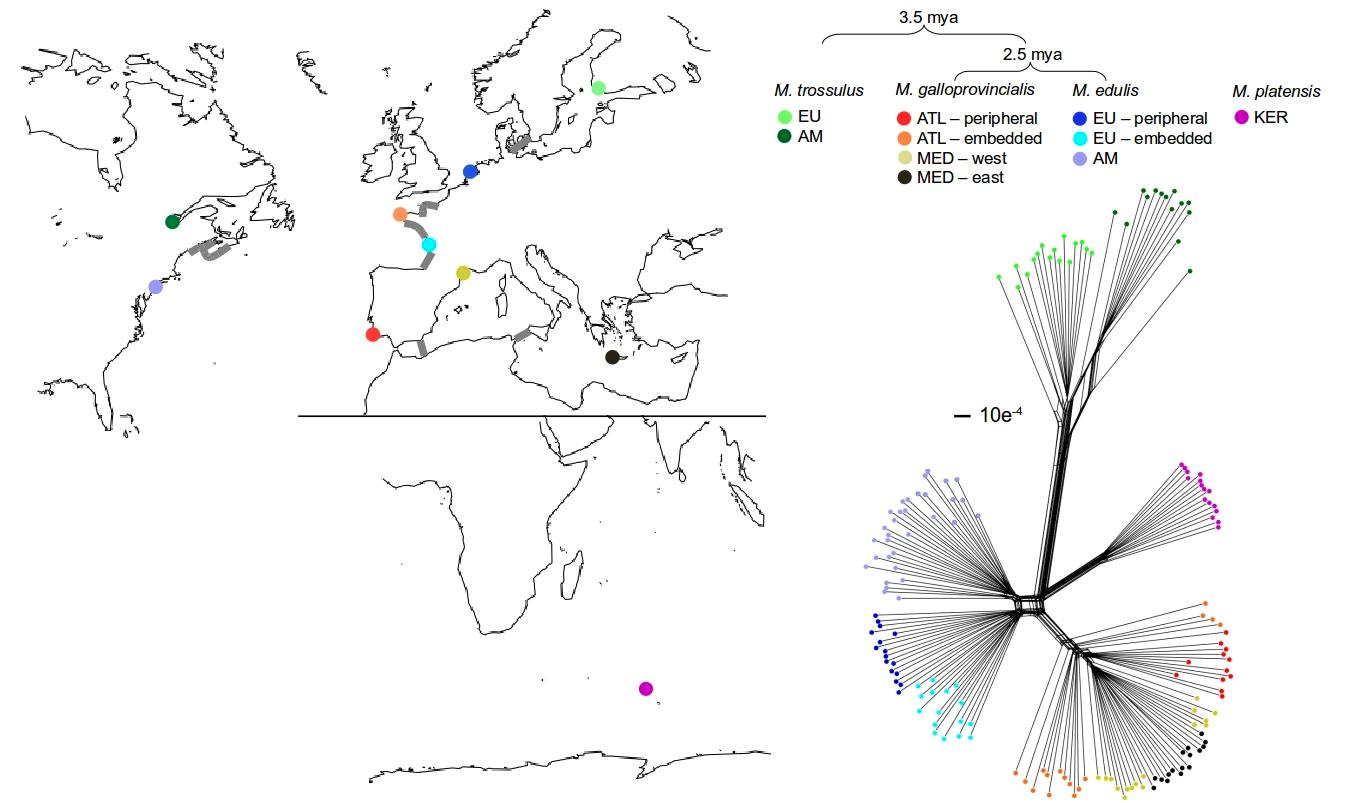
Localities and genetic network of the Northern and Southern *Mytilus* samples (ten GBS-typed populations). The genetic network was produced with the neighbour-net method based on 51,878 high-quality GBS SNPs. The *Mytilus platensis* sample is ‘Baie de la Mouche’ (KER, purple) in the Kerguelen Islands (Southern Ocean). *Mytilus trossulus* samples are ‘Tvarminne’ (EU, light green) in the European population of the Baltic Sea and ‘Tadoussac’ (AM, dark green) in the American population of the Saint Lawrence River. *Mytilus galloprovincialis* samples are ‘Faro’ (ATL – peripheral, red) in the Atlantic population of Iberian Coast, ‘Guillec’ (ATL – embedded, orange) in the Atlantic population of Brittany, ‘Sete’ (MED – west, yellow) in the Occidental Mediterranean basin and ‘Crete’ (MED – east, black) in the Oriental Mediterranean basin. *Mytilus edulis* samples are ‘Wadden Sea’ (EU – peripheral, light blue) in the European population of the North Sea, ‘Lupin/Fouras’ (EU – embedded, cyan) in the European population of the Bay of Biscay and ‘Quonochontaug/Old Saybrook Town’ (AM, slate blue) in the American population of Rhode Island. Estimate of divergence times between Northern species are indicated with braces (Rawson & Hilbish 1995, Roux *et al.* 2014), and contact zones between them are depicted by grey stripes.

### KASPar SNP panel

Based on the SNP database generated by GBS, we specifically selected SNPs segregating in the eight GBS-typed Kerguelen individuals to analyse the fine-scale genetic structure in the island, and its relation to the local environment. Moreover, as we wanted to determine if the micro-geographic structure in the Kerguelen was primarily driven by standing variation in the Northern complex of species (i.e. SNPs fixed between Northern species), the selected SNPs were not a random sample of the SNPs detected by GBS, otherwise they would have been mainly private polymorphisms to the Kerguelen (60% of the Kerguelen SNPs were found private). As such, we further enriched our SNP array with ancestry-informative markers, the most differentiated SNPs between pairs of Northern Hemisphere species (representing 33% of the non-private Kerguelen SNPs, 10% of the whole SNP dataset), namely the West-Mediterranean *M. galloprovincialis* population, the North-Sea *M. edulis* population and the Baltic-Sea *M. trossulus* population. FST values (Weir & Cockerham 1984) were calculated using the R package hierfstat (Goudet 2005) for each SNP between pairs of populations (Text S1). SNPs in the upper 15% of the empirical FST distribution were categorized as highly-differentiated. SNPs with more than 25% of missing data were discarded. SNPs were further filtered-out based on Illumina Assay Design Tool scores (available on Illumina web page, http://support.illumina.com) which predicts probes success based on the number of degenerated sites in the flanking sequences (250 bp on each side of the focal SNP). The final array comprised 58 SNPs out of which 30 were highly differentiated between Northern species (11 *M. trossulus*-specific, 8 *M. edulis*-specific and 10 *M. galloprovincialis*-specific, Table S2).

### KASPar genotyping in the Kerguelen Islands

We used samples collected from 35 sites in the Kerguelen Islands by Gérard *et al.* (2015), totalling 695 individuals (Supp. Info. M&M and Table S3). We additionally included other samples from the Southern Hemisphere (Gérard *et al.* 2008) to assess the genetic relationships previously proposed by Borsa *et al.* (2012) with a handful of markers: one sample from Western Australia in Nedlands (AUS, n=12 individuals); two samples from New Zealand, Dunedin (DUN, n=8) and Wellington Harbour (WHL, n=10); two samples from Tasmania, the Simpson’s Bay (SIM, n=8) and Hobart (HOB, n=9); one sample from Chile in Maullin (MAU, n=15). Pieces of mantle tissue were preserved in 95% ethanol, and DNA was extracted with the Macherey-Nagel NucleoSPin 96 Tissue kit. We used KASPar (Kompetitive Allele Specific PCR, Smith & Maughan 2015), a fluorescence-based genotyping assay of allele-specific PCR products, to genotype the 58 SNPs, of them, 44 SNPs were successfully amplified. We removed seven loci which showed significant FST values between the eight GBS-typed Kerguelen individuals and the KASPar individuals. These may be due to error in the genotyping-by-sequencing, typically the assembly of paralogous loci in two alleles of the same locus, or alternatively to problem of amplification in the KASPar assay as a consequence of primer design. More generally, these discrepancies call for studies that quantitatively compare the two genotyping methods. We further eliminated two loci with null alleles (significant FIS values in most of the sampling sites) and two loci physically linked to one another. The final dataset was composed of 33 KASPar SNPs (Table S2). Additionally, we included allele frequency data of a length-polymorphism locus in the adhesive plaque protein gene, Glu-5’, previously scored in the same sampling sites (Gérard *et al.* 2015). Genotypes for all individuals at each KASPar SNP is available in Text S2, and population allele frequencies are given in Table S4.

### Genetic network of the *Mytilus* spp

Genotypes of the GBS dataset were statistically phased with beagle v3.3.2 (Browning &Browning 2007) using genotype likelihoods provided by bcftools. All individuals were included in the analysis to maximize linkage disequilibrium, and 20 haplotype pairs were sampled for each individual during each iteration of the phasing algorithm to increase accuracy. Phased sequences (haplotypes) were then generated using a custom perl script. An individual genetic network analysis was conducted with splitstree4 v4.12.6 (Hureson & Bryant 2006) to get insight into the population relationships across the three Northern Hemisphere species and the eight individuals sampled in the Kerguelen Islands. All haplotype loci were compiled to create a composite chromosome of 51,878 high-quality SNPs and analysed using the neighbour-net method.

### Analyses of admixture in the *Mytilus* spp

An estimation of the historical relationships among the nine Northern populations and the GBS-typed Kerguelen population was performed with *TreeMix* v.1.1 (Pickrell & Pritchard 2012). TreeMix jointly estimates population splits and subsequent admixture events based on the F-statistics introduced by Reich *et al.* (2009), and commonly recognised as a valid support for admixture (e.g., in humans: Pickrell & Pritchard 2012, Wong *et al.* 2017). A maximum-likelihood population tree was estimated based on the matrix of GBS allele frequency covariance between population pairs, and admixture events were sequentially added. To account for linkage disequilibrium, variants were grouped together in windows of size k=100 SNPs. Trees were rooted with the two *M. trossulus* populations and no sample size correction (option “-noss”) was applied. We tested for a range of migration events from m=0 to m=12, and looked for an asymptotic value of the log-likelihood. The number of significant migration events was assessed by stepwise comparison of Akaike information criterion. Finally, we made 100 bootstrap replicates (option “– bootstrap”) of the maximum-likelihood tree to assess statistical support of migration events.

Based on the GBS genotypes, we additionally performed a model-based clustering analysis of these populations, which assumes admixed individuals with contributions from *K* panmictic reference populations. Ancestry of each individual was estimated using the maximum-likelihood approach implemented in ADMIXTURE v1.23 (Alexander *et al.* 2009). We ran 50 replicates for a number of clusters from K=2 to K=8 and chose the maximum log-likelihood run for each K. We also performed a supervised ADMIXTURE analysis (in which the reference populations were provided) on the Kerguelen individuals with the KASPar SNPs (K=2 clusters and 50 replicates). We defined *M. edulis* and the *M. platensis* Chilean mussels as ancestral populations from which the Kerguelen individuals derive their ancestry. Individual ancestries are provided in Text S3 for the GBS analysis and Text S4 for the KASPar analysis.

In complement to the *TreeMix* analysis, we used a model-based approach implemented in ∂a∂i v1.6.3 (Gutenkunst *et al.* 2009) to explicitly test for the presence of gene flow. This method has proven useful for distinguishing ancestral shared polymorphism between recently-diverged species evolving under strict isolation from a scenario including migration between them (e.g., in *Arabidopsis*: Hubert *et al.* 2014; sea bass: Tine *et al.* 2014; poplars: Christe *et al.* 2017; whitefish:Rougeux *et al.* 2017). We assessed the divergence history between Kerguelen mussels and Northern species in a pairwise manner based on their folded joint site frequency spectrum at the cDNA contigs (provided in Text S5 in ∂a∂i format, and plotted in Figure S1): Kerguelen vs. *M. edulis* (represented by the European sample “EU – peripheral”); Kerguelen vs. *M. galloprovincialis* (Mediterranean sample “MED – west”); Kerguelen vs. *M. trossulus* (European sample “EU”). We defined fourteen demographic models following previous studies (Tine *et al.* 2014; Christe *et al.* 2017, see Figure S1) to test: (i) the timing of gene flow during divergence (absence of gene flow “SI”, continuous migration “IM”, secondary contact “SC”, ancient migration “AM”, periodic secondary contact “PSC” with two periods of current gene flow, periodic ancient migration “PAM” with two periods of ancient migration, and a three-period model of ancient migration, isolation and current gene flow “AMSISC”); (ii) the genomic heterogeneity in gene flow (presence “2M” or absence of interspecific genomic barriers); (iii) the genomic heterogeneity in effective population size (presence “2N” or absence of Hill-Robertson effects). All models began with the split of the ancestral population in two daughter populations, and then were followed by divergence in the absence or presence of gene flow. Each model was fitted to the observed joint site frequency spectrum (singletons were masked) using three successive optimization steps: “hot” simulated annealing, “cold” simulated annealing and BFGS-Broyden–Fletcher–Goldfarb–Shanno-algorithm (Tine *et al.* 2014). Model comparisons were made using Akaike information criterion. A summary of the models is given in Table S5 and the script that defines the models in ∂a∂i is given in Text S6.

### Topology weighting of the *Mytilus* spp

The distinct haplotype loci of the GBS dataset were also individually analysed with the neighbour-net method. Allele genealogies were inferred with the R package APE (Paradis 2010) using a neighbour-joining algorithm with F84 distances (Felsenstein & Churchill 1996). Haplotype loci were filtered based on the following excluding criteria: scale < 0.00005; 0.00005 =< scale < 0.0005 & length < 10000 bp; 0.0005 =< scale < 0.001 & length < 5000 bp; and scale >= 0.001 & length >= 1000 bp, where “scale” is the scale of the gene tree and “length” is the length of the sequence. Neighbour-joining trees of the 395 retained sequences are available in Text S7 and their length are indicated in Table S6 (4.5 kb in average, a minimum length of 1 kb and a maximum length of 25 kb).

For each haplotype locus, the relationships between the Northern species and the Kerguelen population were then quantified using *Twisst*, a tree weighting approach that has been successfully applied to *Heliconius butterflies* to evaluate the support of different phylogenies around colour pattern loci (Van Belleghem *et al.* 2017). We tested the three possible unrooted topologies: (A) *M. edulis* grouped with the Kerguelen population; (B) *M. galloprovincialis* grouped with the Kerguelen population and (C) *M. trossulus* grouped with the Kerguelen population. Their exact weightings to the full tree were estimated by considering all subtrees (“complete method”). Only contigs with a resolved topology were analysed: 67 contigs for which one topology had a weight greater or equal to 0.75. These topologies were further classified in two categories depending on whether they most plausibly reflect: (i) ancient divergence of the Kerguelen population (i.e. the Kerguelen and Northern individuals clustered into distinct monophyletic groups) or, (ii) introgression with one of the Northern species (i.e., the Kerguelen individuals were distributed within one or more Northern clades); "na" stands for topologies that we were unable to classify in these two categories due to a lack of informative sites. Tree topology weightings and classification are available in Table S6.

### Analyses of genetic variation in the Kerguelen Islands

For each KASPar SNP, estimation of F_ST_ values (Weir & Clark Cockerham 1984) was calculated over all sampling sites (Table S2), and in a pairwise manner across all SNPs (Table S7) using Genetix 4.05 (Belkhir *et al.* 2002). Their significance was tested by a permutation procedure (1000 permutations) and adjusted with the Bonferroni’s correction for multiple comparisons (Benjamini & Hochberg 2000).

### Analysis of habitat variables in the Kerguelen Islands

To evaluate how much of the genetic variation among sites was explained by local environmental factors, we used redundancy analysis (RDA), a constrained ordination method implemented in the R package vegan (Oksanen *et al.* 2017). It performs a multiple linear regression between a matrix of response variables (individual genotypic data) and a matrix of explanatory variables (environmental factors). Notably, the effect of partially confounded explanatory variables can be estimated separately. RDA is commonly used to estimate the relative contribution of spatial and environmental components on species communities, and it has been recently applied to analysis of population genetic structure (e.g., Legendre & Fortin 2010). Geographic coordinates and five qualitative factors were measured in each site to describe the local habitat (Table S3): (i) Substrate (rock: R, blocks: B, gravels: G, or sand: S); (ii) Wave Exposure (sheltered: Sh, or exposed: E); (iii) Slope (flat: F, steep: St, or hangover: H); (iv) Salinity (oceanic water: OW, or low-salinity water: LSW); (v) *Macrocystis* (presence: P, or absence: A).

We specifically tested the effect of each of these constrained factors (explanatory variables) on the distribution of genotypes at the 33 KASPar SNPs (response variables). The following initial model was used: Genotypes ~ *Macrocystis* + Salinity + Slope + Exposure + Substrate + Longitude + Latitude. The significance of the global model was first established by permutation test, in which the genotypic data were permuted randomly and the model was refitted (1000 permutations). Marginal effect permutation tests were then performed to assess the significance of each factor by removing each term one by one from the model containing all other terms (1000 permutations). Non-significant factors were removed from the final model. Based on that simplified model, we performed a conditioned RDA analysis for each factor to assess its contribution to the genotypic variance independently from the other explanatory variables. These co-variables were removed from the ordination by using a condition function: Genotypes ~ tested variable + condition(all other variables). Finally, we performed a conditioned RDA on geography to specifically control its confounding effect: Genotypes ~ significant environmental variables + condition(significant geographic variables).

### Simulations of secondary contact

Following the methodology of Gagnaire *et al.* 2015, we modelled a secondary contact between two semi-isolated genetic backgrounds (*M. edulis* vs. *M. platensis* Chilean population) that meet twice on a circular stepping stone model of 30 demes (between demes n°18 / n°19 and demes n°19 / n°20), and start to exchange genes. At generation zero, the *M. edulis* background settles in deme n°19 while the *M. platensis* background is located everywhere else. Their initial allele frequency at the barrier loci was set-up to 0.10 for the *M. edulis* background and 0.70 for the *M. platensis* background, which correspond to the average foreign allele frequency at the four most differentiated loci observed in the *M. edulis* and Chilean *M. platensis* mussels, respectively (see Figure 4B). The auto-recruitment rate was set to 1-*m*, and migration to adjacent demes was *m*/2 (with *m*=0.5). A barrier to dispersal was set between demes n°18 and n°19 (*m*=0.05), which corresponds to the genetic break observed between sites PAF and RdA in the Kerguelen Islands. Selection acts in haploid individuals on a two-locus incompatibility, which is linked to a neutral marker located 1cM away and unlinked to a second neutral marker. Strong and asymmetric selection were required to maintain the two genetic backgrounds after contact (s=0.5 against the *M. platensis* allele in the *M. edulis* background vs. s=0.2 against the *M. edulis* allele in the *M. platensis* background). Deme size was constant and set to 500 individuals.

## Results

### The Kerguelen mussels: signal of divergence of a Southern lineage after transoceanic migration and admixture with Northern lineages

An individual genetic network (Figure 1) was built from a subset of 51,878 high-quality SNPs genotyped in nine Northern populations and eight individuals from the Kerguelen Islands. We observed that the Northern populations formed three distinct clusters, corresponding to the three Northern species: *M. edulis*, *M. galloprovincialis* and *M. trossulus*. Accordingly, the majority of SNPs fixed between populations (295 in total) were species-specific: *M. edulis*=6, *M. galloprovincialis=*62 and *M. trossulus*=224. The Kerguelen individuals clustered together into a single clade. Indeed, the proportion of SNPs which were private to the Kerguelen Islands amounted to 60% (3,805 private for a total of 6,297 SNPs in Kerguelen, after removing singletons). In comparison, the number of private SNPs in *M. trossulus* was 3,070, and it was only 492 in *M. galloprovincialis* and 48 in *M. edulis* (indicative of introgression between the two latter species). Among the 2,492 SNPs shared by the Kerguelen mussels with Northern species, 33% (830) were highly differentiated between at least two Northern species. When considering Northern species-specific SNPs, 83% of those fixed in *M. edulis* were segregating in the Kerguelen (5 for a total of 6 fixed). These numbers were 16% for *M. galloprovincialis* (10 for a total of 62 fixed) and 12% in *M. trossulus* (27 for a total of 224 fixed). A multivariate analysis on KASpar-typed SNPs, including the Northern samples, the Kerguelen Islands and other samples from the Southern Hemisphere that were also genotyped in our SNP assay, is provided as a supplementary figure (Figure S2). The principal component analysis clearly shows that the Chilean mussels (MAU) group with the Kerguelen mussels in accordance with them being both named *M. platensis*; while the Australasian samples (Australia, Tasmania and New-Zealand), usually named *M. planulatus*, cluster with the Northern *M. galloprovincialis*. These findings corroborate previous results based on mitochondrial DNA (Gérard *et al.* 2008) and nuclear markers (Borsa *et al.* 2012).

The species relationships found in the genetic network (Figure 1) were generally supported by the maximum-likelihood population tree inferred by *TreeMix* (Figure 2A), except that the Kerguelen population was inferred as the sister-group of *M. edulis*. The pairwise population residuals in a model without admixture (Figure S3) suggested substantial migration between species. So, we sequentially allowed from 0 to 12 migration events in the analysis, and assessed their significance by stepwise comparison of Akaike information criterion (Figure S3). The best fit to the data was obtained with seven migration events, which significantly improved the log-likelihood of the model (Figure S3). This population tree was bootstrapped 100 times to assess statistical support of migration events. They were generally weakly supported, with only four migration events that had more than 50% bootstrap support (Figure 2A and Table S8). The most robustly inferred migration event was between the Mediterranean *M. galloprovincialis* and the Kerguelen population (81 % of bootstrap replicates). The two others included migration among Northern species as expected: the European populations of *M. edulis* and *M. galloprovincialis* (51%), and the European populations of *M. edulis* and *M. trossulus* (68%). Migration events were also inferred between the Mediterranean *M. galloprovincialis* and the European *M. trossulus* (55%), as well as between the Kerguelen population and the American *M. trossulus* (38%). Reticulated evolution with the American *M. trossulus* was further supported by their excess of privately shared polymorphism with the Kerguelen population (Figure 2B). Migration between the Kerguelen population and the European *M. edulis* was detected, but only in 4% of bootstrap replicates.

**Figure 2.**
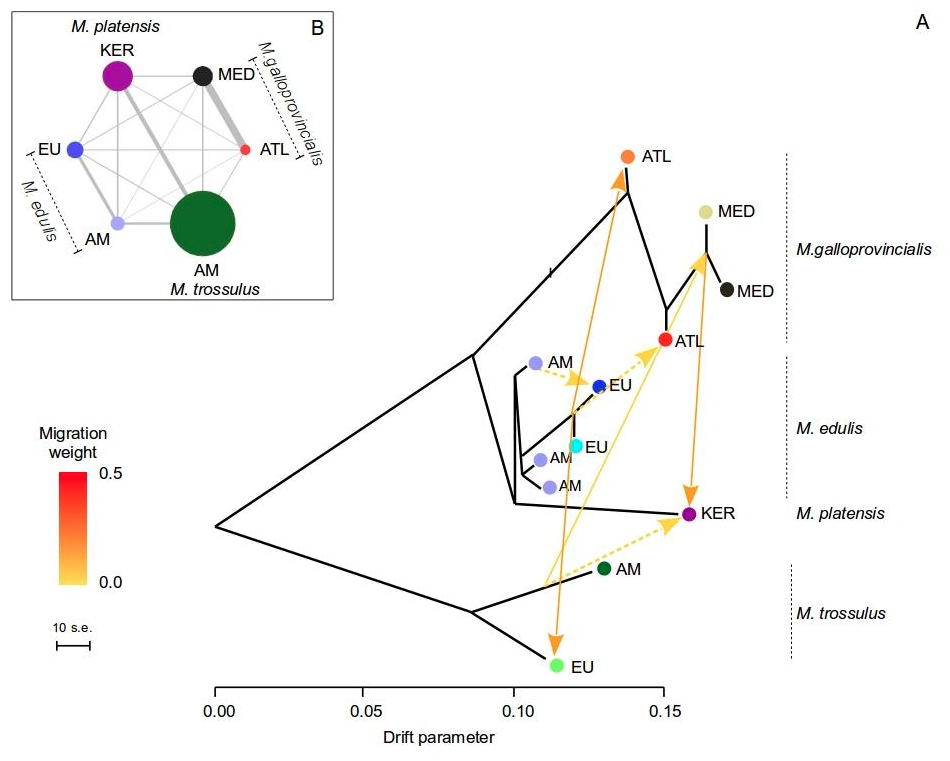
Evidence of admixture in the ten GBS-typed *Mytilus* spp. populations. **A.** Maximum-likelihood population tree inferred by *TreeMix* showing 7 migration events. ML estimation was made by using 32,162 GBS SNPs grouped in windows of 100 SNPs. Terminal nodes are labelled by locality abbreviation (Figure 1 and Table S1) and colors match Figure 1. The drift parameter is shown on the x-axis. Horizontal branch lengths are proportional to the amount of genetic drift that has occurred in each branch. Admixture arrows are coloured according to the migration weight, which represents the fraction of ancestry derived from the migration event. Migration events with bootstrap support less than 50% are shown with dotted lines, and those with bootstrap support less than 20% are not shown (Table S8). **B.** Pairwise privately shared SNPs between a subset of populations of similar sample size (Table S1). The width of the lines is proportional to the number of SNPs shared (*M. trossulus:* KER-AM=1,394 (America); *M. galloprovincialis*: KER-ATL=254 (Atlantic - peripheral) and KER-MED=392 (Occidental Mediterranean basin); *M. edulis*: KER-EU=421 (Europe - peripheral) and KER-AM=532 (America)). The size of the circles is proportional to the number of non-shared private SNPs (KER=6,022; *M. trossulus*: AM=13,095; *M. galloprovincialis*: ATL=1,948, MED=3,927; *M. edulis*: EU=3,251, AM=2,731). Singletons were removed.

Our reconstruction of the divergence history with ∂a∂i consolidated evidence for admixture between the Kerguelen mussels and each Northern species, although pointed out to a periodic connectivity with several periods of admixture. In all three comparisons, the models without gene flow were consistently the least supported (“Strict Isolation”, Table S5). In the *M. edulis* and *M. galloprovincialis* comparisons, the model of periodic ancient migration with varying rates of introgression among loci (“*PAM_2M*”) received the strongest statistical support (Table S5).

Migration occurred right after the split as well as later in the divergence during a relatively short period (~15% to 25% of the total divergence time) and it was asymmetric with a substantial fraction (85% to 90%) of the mussel genome in the Kerguelen permeable to *M. edulis* and *M. galloprovincialis* gene flow. In the *M. trossulus* comparison, the three-period model with homogeneous rates of introgression among loci (“AMSISC”) was the best supported (Table S5). A first period of ancient migration lasted ~85% of the divergence time, and was followed by a short period of isolation and a very short secondary contact (~2% of the total divergence time). Kerguelen mussels were mostly resistant to *M. trossulus* introgression (Table S5). Overall, these results suggest that the Kerguelen mussel is a Southern lineage related to *M. edulis* and that it admixed with all three Northern species (*M. edulis*, *M. galloprovincialis* and to a lesser extent with *M. trossulus*) in the past, either after secondary contact and/or early in the divergence of the species.

### Variation of admixture histories across the genome

To further investigate how genetic relationships varied across the genome, we quantified the contribution of three unrooted topologies (Figure 3) to the full tree at 395 GBS contigs with *Twisst*. Only 17% (67) of them showed resolved relationships, i.e. one of the unrooted topology weighted 75% or more, among which 40% (27) were highly resolved (weight >= 90%). The reason for this low rate of resolved relationship is twofold: (i) ongoing sorting of shared ancestral polymorphism, and (ii) introgression. In the Northern Hemisphere introgression has been shown to be the main process thanks to model fitting (Fraïsse *et al.* 2018) and using comparison of introgression differential between populations within species (Fraïsse *et al.* 2016). It is more difficult to tell apart ancestral shared polymorphism from introgression for the single Kerguelen population. Above analyses suggest introgression is contributing although it is difficult to evaluate its contribution. The most represented resolved topology (39 contigs) put the Kerguelen individuals together with *M. edulis*, while they were grouped with *M. trossulus* in 19 contigs (i.e. ancestral to the *M. edulis* / *M. galloprovincialis* subgroup) and with *M. galloprovincialis* in 9 contigs (Table 1). This illustrates a high rate of ancient incomplete lineage sorting that resolved into alternative topologies after the consecutive speciation events, which resulted in the three species *M. edulis*, *M. galloprovincialis* and *M. platensis*. In other words, the transequatorial migration that leads to *M. platensis* occurred soon after the split between *M. edulis* and *M. galloprovincialis*.

**Figure 3.**
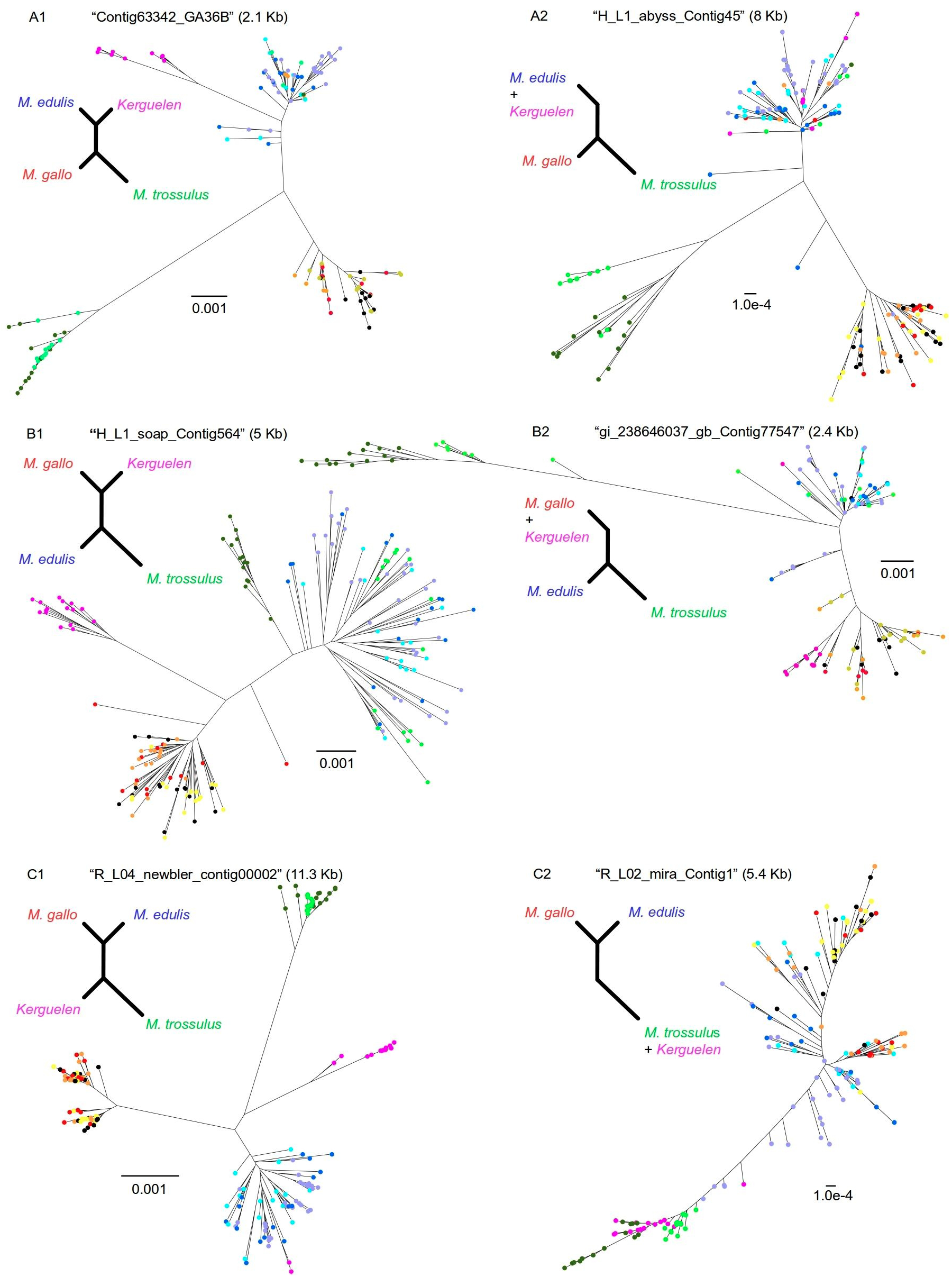
Summary of the different gene topologies obtained with *Twisst* for the GBS dataset. Three topologies have been weighed with *Twisst* for each of the 395 contigs (of which only 67 were topologically resolved), and classified in different categories depending whether the Kerguelen mussels branched as a sister-clade to a Northern species (“ancient divergence”), or were distributed within a Northern species (“introgression”). **A.** Kerguelen clustered with *Mytilus trossulus*: A1 “ancient Kerguelen divergence” and A2: “introgression”; **B.** Kerguelen clustered with *Mytilus edulis*: B1 “ancient Kerguelen divergence” and B2: “introgression”; **C.** Kerguelen clustered with *Mytilus galloprovincialis*: C1 “ancient Kerguelen divergence” and C2: “introgression”. For each category, a typical neighbour-joining tree computed on the longest non-recombining block of the contig is shown (defined with the Difference of Sums of Squares method of McGuire & Wright 2000). In panel A2, the *Mytilus edulis* haplotypes have totally replaced their Southern counterparts in the Kerguelen. Panel B2 suggests a more ancient introgression of *Mytilus galloprovincialis* haplotypes given that all haplotypes sampled in the Kerguelen form a distinct cluster within the *Mytilus galloprovincialis* clade. And panel C2 represents a complete introgression of *Mytilus trossulus* haplotypes into the Kerguelen. Contig IDs correspond to Table S6, and sizes of non-recombining blocks are given in brackets. Colors match Figure 1.

**Table 1.** Counting the different gene topologies for the GBS dataset (10 populations, 67 gene trees)

| topology | categories | counts |  |
| --- | --- | --- | --- |
| | | weight $\geq 0.90$ | weight $\geq 0.75$ |
| *A. M. edulis* with Kerguelen | A1 ancient Kerguelen divergence | 3 | 5 |
|  | A2 introgression | 5 | 12† |
|  | NA | 5 | 22 |
|  | **total** | **13** | **39** |
| *B. M. galloprovincialis* with Kerguelen | B1 ancient Kerguelen divergence | 2 | 2 |
|  | B2 introgression | 3 | 3 |
|  | NA | 0 | 4 |
|  | **total** | **5** | **9** |
| *C. M. trossulus* with Kerguelen | C1 ancient Kerguelen divergence | 5 | 11 |
|  | C2 introgression | 3 | 4 |
|  | NA | 1 | 4 |
|  | **total** | **9** | **19** |
| **TOTAL** |  | **27** | **67** |

**topology**: three topologies have been weighted with *Twisst* (among the 395 gene trees, only 67 were topologically resolved); **categories**: “*ancient divergence*”: the Kerguelen population branched as a sister-clade to a Northern species; “*introgression*”: Kerguelen mussels were distributed within a Northern species; “*na*”: topologies that we were unable to classify in these two categories. **weight**: exact weightings of each topology to the full tree.

When classifying the topologies in categories, 18 contigs supported the “ancient Kerguelen divergence” scenario while 19 supported an “introgression” scenario among which 4 were from *M. trossulus*, 12 from *M. edulis* and 3 from *M. galloprovincialis*; 30 contigs could not be classified. It should be noted that all these patterns hold when using a minimal weight of 90% (Table 1). Figure 3 illustrates representative cases of the different *Twisst* categories, including candidate loci for introgression (A2, B2 and C2 in Figure 3). These results suggest that the Kerguelen mussels have a genome of mixed ancestry, mainly dominated by *M. edulis,* from which they are closely-related, but with which they also have probably secondarily admixed. This is in accordance with the *TreeMix* analysis where the Kerguelen population was inferred to be the sister-clade of *M. edulis* and experienced admixture events with *M. edulis* and *M. galloprovincialis*. However, this seems quantitatively different from the ∂a∂i results in which both *M. edulis* and *M. galloprovincialis* broadly contributed to the genomic composition of the Kerguelen mussels. However, we call for caution when comparing these two methods as ∂a∂i provides genome-wide migration rate estimates, while *Twisst* relies on counting different types of topologies across the genome and many gene trees could not be classified, and *TreeMix* infer single admixture event under the hypothesis of homogeneous introgression (while *Mytilus* species barriers are often semi-permeable, Roux *et al.* 2014, Fraïsse *et al.* 2018, and see Table S5).

### Substantial genetic structure in the Kerguelen Islands

Mussels were collected from 35 sampling sites all around the Kerguelen Islands (Figure 4A, Table S3) and successfully genotyped at 33 KASpar SNPs. Pairwise FST values across all SNPs (Table S7) revealed significant fine-scale genetic differentiation between sites from different geographic regions. Remarkably, RdA (North-East) and PCu (West) were significantly differentiated with nearly all other sites. Sites from the South, especially BdS, and from the North, especially AS, were differentiated from the Gulf of Morbihan. At a smaller scale within the Gulf of Morbihan, genetic structure among several sites was detected, but their significance level did not pass the correction for multiple tests. These results extend the study by Gérard *et al.* (2015) to many SNPs and substantiate their finding of significant genetic differentiation at different scales in the island.

**Figure 4.**
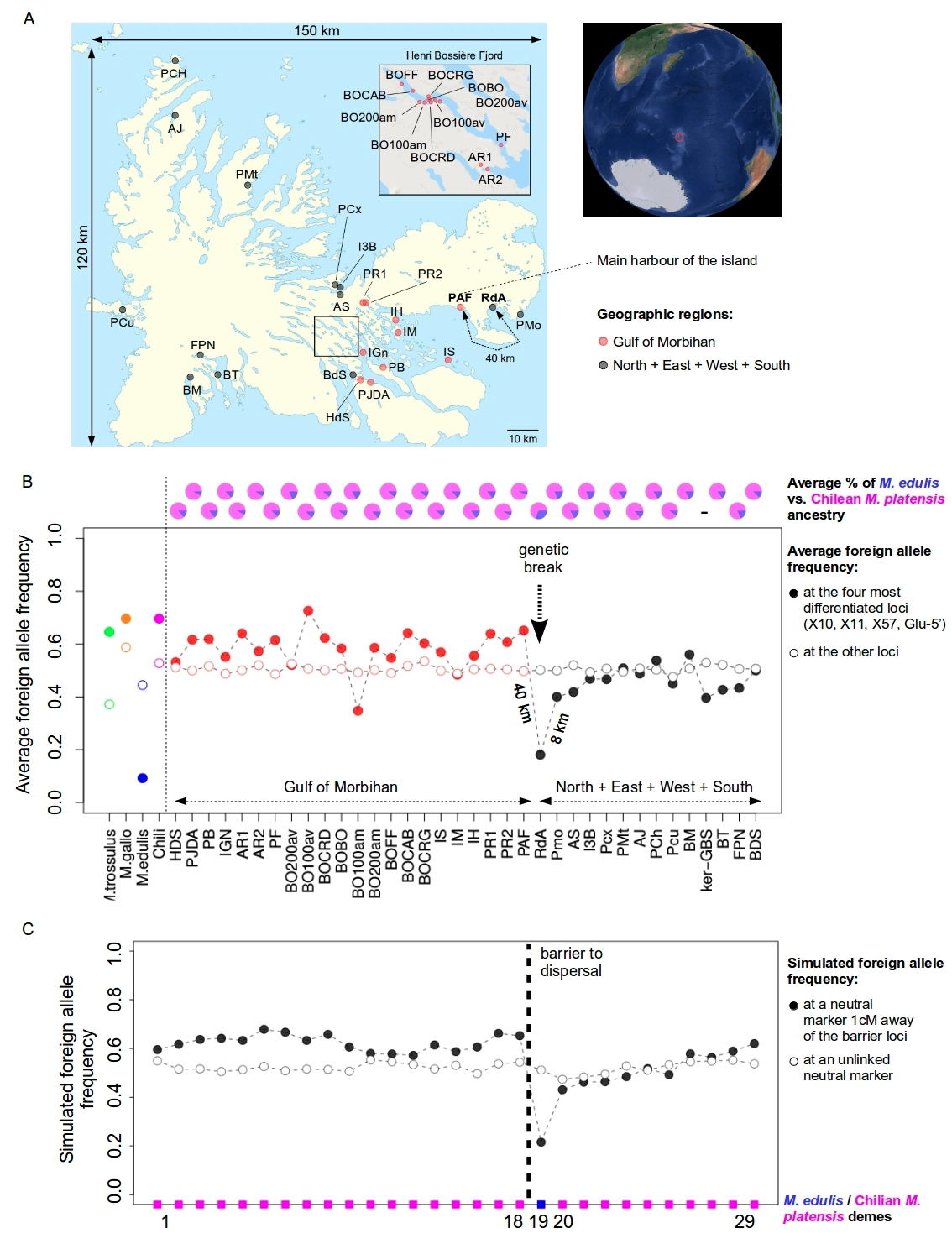
Geographic variation of the average foreign allele frequency across the four most differentiated loci in the Kerguelen Islands (filled symbols, X10, X11, X57 and Glu-5’ from Gérard *et al. (*2015)) and the other loci (open symbols). Alleles were labelled based on their frequencies in the *M. galloprovincialis* Atlantic population of Iberian Coast (Table S4). Red points indicate sites located in the Gulf of Morbihan. **A.** Map of the Kerguelen Islands (150 km East to West; 120 km North to South) together with a world map indicating its location in the Southern Ocean (surrounded in red), and an enlarged map of the Henri Bossière Fjord. Sites PAF (the main harbour of the island) and RdA are separated by 40 kms of coasts, RdA and Pmo by 8 kms, and they show a genetic break on Panel B. Sampling details are provided in Table S3. **B.** Frequency of the average foreign allele across sampling sites in the Kerguelen Islands (ordered by geography), in the three reference Northern species and in the Chilean mussels. See Figure S4 for the detailed pattern at each locus. On top, the average *Mytilus edulis* vs. Chilean ancestries from an ADMIXTURE analysis (K=2) are shown for each site. **C.** Simulation results after 300 generations of a secondary contact between *Mytilus edulis* and Chilean *Mytilus platensis* mussels that meet twice on a circular stepping-stone model (between demes n°18 / n°19 and demes n°19 / n°20). A physical barrier to dispersal was modelled between the *Mytilus edulis* deme (n°19) and the Chilean deme n°18 (dashed vertical bar, migration rate=0.05 instead of 0.5 as everywhere else). Deme size was fixed to 500 individuals, and the species barrier was asymmetric (at barrier loci, the selection coefficient against *Mytilus edulis* in the *Mytilus platensis* background was set to 0.2 while it was set to 0.5 against the *Mytilus platensis* allele in the *Mytilus edulis* background).

Global FST across all sites was calculated for each SNP and tested with 1000 permutations (Table S2). Values were non-significant after Bonferroni’s correction, except at the three most differentiated loci, which are ancestry-informative: X10, X11 and X57. Their foreign allele, defined based on its frequency in the Northern species (*M. galloprovincialis* Atlantic population of Iberian Coast, Table S4), was at low frequency in the North of the island, especially in the North-East sites, RdA and PMo. In contrast, it was at higher frequency in the Gulf of Morbihan and at intermediate to low frequency in the South and West. These trends were similar to those at Glu-5’ (Table S4), a nuclear marker suspected to be affected by differential selection in the island (Gérard *et al.* 2015) and at candidate allozymes although with fewer sampling sites (Blot *et al.* 1989). Across all sites, the frequencies of the foreign allele at Glu-5’ were significantly correlated with those at X10 (r=0.61, p-value < 0.001), X11 (r=0.419, p-value=0.012), and X57 (r=0.49, p-value=0.003), but they were globally higher at Glu-5’ (Figure S4).

The foreign allele frequency at those four loci is represented in Figure S4 and the average over the four loci in Figure 4B (filled symbols). These clearly show a genetic break between two geographically close sites, PAF and RdA (40 kms apart), and to a lesser extent between RdA and Pmo, which are separated by only 8 kms of coasts. The average frequency was the highest in the Gulf of Morbihan (from HdS to PAF), then it abruptly dropped down (in 40 km) between PAF and RdA (respectively on the West and East coast of the Prince of Wales’ Peninsula), and finally increased gradually along the coast from North-East to South-West. Notably, the variance in allele frequency at the four loci was weak (0.0208 in average across localities). This is in sharp contrast with the pattern observed at the other loci (open symbols) of which the average frequency remained similar across all sites (Figure 4B), and variance was stronger (0.0938 in average across localities).

An admixture analysis using all KASPar SNPs and defining *M. edulis* and the Chilean mussels as reference populations (Figure 4B, pie charts), suggests that the Kerguelen Island is occupied by mussels related to Chilean mussels (*M. platensis*), and that RdA has by far the highest level of *M. edulis* ancestry (31% compared to <19% elsewhere). We therefore hypothesize that two genetic backgrounds may be present in the island, one related to *M. edulis* and trapped at site RdA close to a potential density trough in the the Prince of Wales’ Peninsula, and the other related to Chilean mussels and present everywhere else. This pattern is theoretically expected (Barton 1979, Barton & Hewitt 1985) as areas of low density or low carrying capacity tend to trap “tension zones” between incompatible genetic backgrounds (similarly to ecological boundaries).

We illustrated this scenario by simulating a secondary contact between these two backgrounds (including a physical barrier to dispersal between demes n°18 and n°19), and tracking the frequency of the foreign allele at two neutral markers (linked and unlinked to the barrier locus) a few hundreds generations after the contact (Figure 4C). Simulations often fitted well with the observed variation in allele frequency across sites, as predicted by Gagnaire *et al.*’s model (2015), provided that the species barrier was asymmetric in order to protect the small *M. edulis* patch to be quickly swamped by *M. platensis* introgression. This suggests that the genetic break at the boundary of the Gulf of Morbihan and the North-East region is better revealed by the frequency of foreign alleles at ancestry-informative loci implying a role of admixture either in the maintenance or in the detection of the genetic structure.

### Environment-associated genetic structure in the Kerguelen Islands

We then tested for genetic-environment associations in the Kerguelen by performing a RDA. Among the seven constrained factors (five qualitative variables, plus geographic coordinates), three were not significant in the initial model (Salinity, Exposure and Latitude, Table S9) and were removed from further analyses. The proportion of total genotypic variance explained by all constrained factors was highly significant in the global model (p-value=0.001, Table 2A, left panel), but quite low (2.32%). The first RDA axis, which explained 61% of the constrained variance, was mainly contributed by *Macrocystis* presence/absence (Figure S5 and Table S10). Accordingly, it was the only factor whose marginal effect remained significant (p-value=0.032, Table 2B, left panel).

**Table 2.** RDA analysis for the KASPar dataset (695 individuals, 33 SNPs)

| RDA (N=695) |  |  |  |  | conditioned RDA (N=695) |  |  |  |  |
| --- | --- | --- | --- | --- | --- | --- | --- | --- | --- |
|  | variance | %variance | d.f. | p-value |  | variance | %variance | d.f. | p-value |
| **A. Global effect** |  |  |  |  | **A. Environmental effect** |  |  |  |  |
| Model | 0.5288 | 2.324 | 7 | 0.001 *** | Substrate+*Macrocystis*+Slope | 0.1213 | 1.0929 | 6 | 0.041 * |
| Residual | 22.2183 | 97.676 | 687 | - | Residual | 10.9621 | 98.7588 | 687 | - |
| **B. Marginal effect** |  |  |  |  | **B. Individual effect** |  |  |  |  |
| Longitude | 0.0128 | 0.0563 | 1 | 0.664 | Longitude | 0.1424 | 0.626 | 1 | 0.001 *** |
| Substrate | 0.0594 | 0.261 | 3 | 0.133 | Substrate | 0.2322 | 1.021 | 3 | 0.001 *** |
| *Macrocystis* | 0.0275 | 0.121 | 1 | 0.032 * | *Macrocystis* | 0.0691 | 0.304 | 1 | 0.011 * |
| Slope | 0.0379 | 0.167 | 2 | 0.217 | Slope | 0.0912 | 0.401 | 2 | 0.072 . |
| Residual | 10.9621 | 48.191 | 687 | - | Residual | 22.2183 | 97.676 | 687 | - |

RDA significance tests are shown on the left for: **(A)** the global model with non-significant terms removed; and **(B)** the marginal effect of each constraining variable, which was tested through permutation tests by removing each term one by one from the model containing all other terms. Conditioned RDA significance tests are shown on the right for: **(A)** the combined effect of environmental variables after conditioning on Longitude; and **(B)** for each term after conditioning on other constraining variables to remove their confounding effects. **N**: number of individual genotypes; **variance**: genotypic variance explained by each factor; **%variance**: percent of genotypic variance; **d.f.**: degrees of freedom; **p-value**: p-values obtained through 1000 permutations (\* $\leq 0.05$ ; \*\* $\leq 0.01$ ; \*\*\* $\leq 0.001$ ).

We statistically controlled for the effect of geography by performing a conditioned RDA analysis on Longitude (Table 2A, right panel). The combined effect of the three environmental variables remained significant (p-value=0.041), explaining 1.1% of the total genotypic variance. Individually, *Macrocystis* presence/absence and Substrate still showed significant effects, after removing all other confounding factors (p-value=0.011 and 0.001, respectively, Table 2B, right panel). This suggests a limited effect of geography on the genetic structure of the mussels, although coastal distance would be a better proxy to account for spatial structure in the island. Interestingly, it has been previously shown that the fine-scale genetic variation at Glu-5’ was also significantly associated with *Macrocystis* presence/absence (Gérard *et al.* 2015). Accordingly, we found a significant correlation between the average foreign allele frequency at the four most differentiated loci in the Kerguelen Islands and the presence/absence of *Macrocystis* (Figure 5A), whereas there was no correlation with the other loci (Figure 5B).

**Figure 5.**
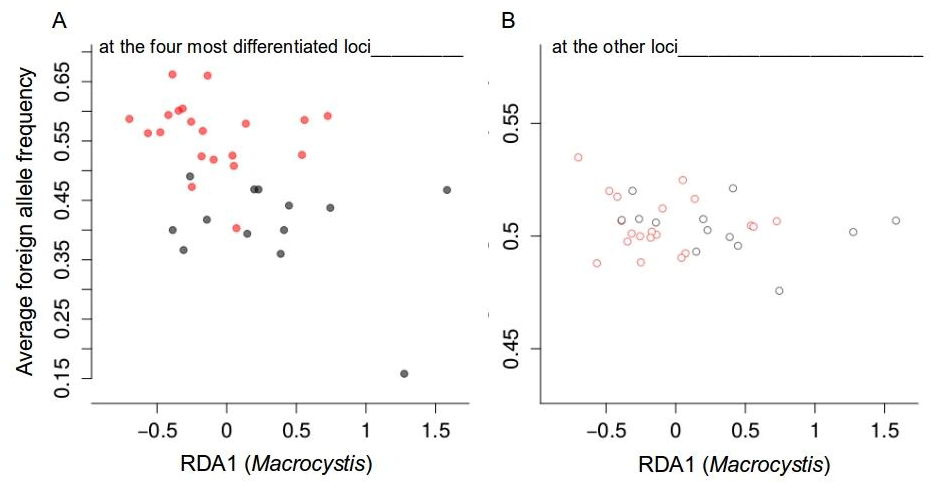
Correlation between the average foreign allele frequency of the four most differentiated loci (**A**) and other loci (**B**) at each sampling site in the Kerguelen Islands (y-axis, Table S4), and the presence/absence of *Macrocystis* (x-axis, this is the average sampling site coordinates on the first axis of the *Macrocystis* RDA, see Table S12). Pearson correlation coefficient: on the left panel, 0.494 (p-value=0.003); on the right panel, 0.107 (p-value=0.541). Red points indicate sites located in the Gulf of Morbihan (see Figure 4A).

The sharp genetic break between RdA and PAF further indicates that two genetic backgrounds may have been locally trapped by an ecological boundary or a region of reduced dispersal (Bierne *et al.* 2011). Accordingly, there is an oceanic threshold at the entrance of the Gulf of Morbihan that impedes exchanges with water masses from outside; and at a larger scale, the Antarctic circumpolar current moves the water masses from West to East causing gyres and turbulences on the North-Eastern coast and pushing water masses far to the East (Karin Gerard, pers. comm.). Thus, the water masses between the Gulf of Morbihan and the North Coast do not mix well, suggesting that exchanges between the two sites are limited. Moreover, these two sites differ at all five ecological variables (Table S3), but not in the direction predicted by their geographical origin: RdA shows an habitat characteristic of the Gulf of Morbihan while being located on the East coast and having the lowest foreign allele frequency (and the reverse is true for PAF). This imperfect association between genotypes and habitats supports the hypothesis that enhanced genetic drift and intense gene flow in the island grambled the signal of genetic-environment association at our markers rather than environmental differential selection.

### Most differentiated SNPs in the Kerguelen Islands are primarily ancestry-informative in the Northern Hemisphere

In the total sample, the average allele frequency of the foreign allele was 0.417 at Glu-5’, 0.503 at X10, 0.619 at X11 and 0.480 at X57. These polymorphisms were well-balanced in the island (i.e. close to 0.5 frequency), despite being species-specific in the Northern species (Table S2, also see Gérard *et al.* 2015 for Glu-5’). To investigate whether micro-geographic structure in the island was primarily depending on Northern ancestry-informative loci, we compared the degree of differentiation between sites in the Kerguelen Islands and that of the Northern species, *M. edulis* and *M. galloprovincialis*, at the 33 KASPar SNPs (Figure 6). Figure 6A shows that the level of genetic differentiation among sites in the Kerguelen (global F_ST_, Table S2) was significantly higher (p-value=0.021) for the ancestry-informative loci (mean=0.015, in orange) compared to the background loci (mean=0.007, in grey). Importantly, the difference between the two categories was also significant when considering the genetic-by-environment association across all variables (Figure 6B: mean_orange=0.112, mean_grey=0.048, p-value=0.006), which was measured by the locus coordinates on the first axis of the conditioned RDA (Table S11); or only including *Macrocystis* presence/absence (Table S12: mean_orange=0.089, mean_grey=0.041, p-value=0.016). Moreover, these patterns hold when adding the locus Glu-5′ (Figure S6) in the case of genetic differentiation (Figure S6A, p-value=0.01), and genetic-environment associations (Figure S6B, p-value=0.004) measured by the FCT from an AMOVA analysis performed by grouping sites according to the presence/absence of *Macrocystis* (see Gérard *et al.* 2015).

**Figure 6.**
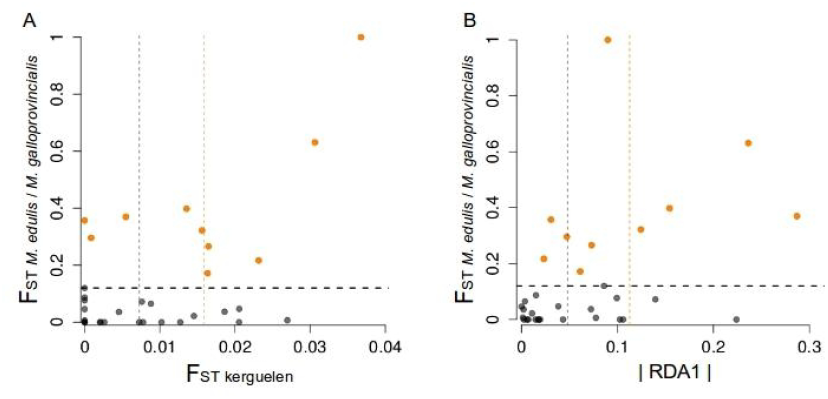
Correlation between the level of differentiation among the Kerguelen Islands (x-axis) and the Northern species (F_ST_, y-axis) at each KASPar SNP. Panel (**A**) shows the genetic differentiation between Kerguelen populations (global F_ST_); and panel (**B**) shows the genetic-by-environment association (locus coordinates on the first axis of the conditioned RDA (absolute values), see Table S11). Northern species are *Mytilus edulis* (EU – peripheral, European population of the North Sea) and *Mytilus galloprovincialis* (MED - west, Mediterranean population of the West basin). Ancestry-informative loci, i.e., F_ST_ *M.edulis_M.galloprovincialis* > 0.120 (horizontal dashed line, see Table S2), are depicted in orange. Wilcoxon’s test between the ancestry-informative loci (orange) and the background loci (grey): (**A**) mean_orange=0.015, mean_grey=0.007 (p-value=0.021); (**B**) mean_orange=0.112, mean_grey=0.048 (p-value=0.006). Their respective means are depicted by vertical dashed lines. Pearson correlation coefficient: (**A**) r=0.540 (p-value=0.001); (**B**) r=0.439 (p-value=0.011).

## Discussion

In this work, we first analyzed the genetic relationship of the Southern Hemisphere Kerguelen mussels with the three Northern species (*M. edulis*, *M. galloprovincialis* and *M. trossulus*) at 1,269 contigs (51,878 SNPs), and we confirmed with this large dataset that mussels in the Kerguelen Islands belong to a Southern lineage (*M. platensis*). We further showed that the Kerguelen population is a sister clade of *M. edulis*.

Our population tree inference provides evidence of genetic exchanges with Northern mussels that occurred after the first establishment in the Southern Hemisphere. This was confirmed by reconstructing their divergence history, which further suggested that introgression occured at variable rates across the genome with some genomic regions resistant to gene-flow while others are essentially permeable. The resulting genome-wide ancestry variation was estimated by applying a new topology weighting method to each GBS sequence (Martin & Van Belleghem 2016), which weighted the contribution of three topologies to the full tree. The majority of the genome showed evidence for both ancient incomplete lineage sorting, which resolved into alternative topologies among the four species studied *(M. trossulus*, *M. edulis*, *M. galloprovincialis* and *M. platensis*), and shared polymorphisms due to ongoing incomplete lineage sorting and introgression among North Hemisphere species and with the Southern lineage, and this resulted in only 17% of the regions with a well-resolved topology. In 51% of these resolved regions, we found clear evidence of admixture, i.e., the Kerguelen haplotypes were all (or part of) nested within a Northern clade. We thus confirmed that the reticulated evolution of the Southern Hemisphere Kerguelen mussels, which was suggested by a handful of nuclear markers (Borsa *et al.* 2007) and mitochondrial DNA (Hilbish *et al.* 2000; Gérard *et al.* 2008), holds at a genome-wide scale. This is in line with recent genomic studies (e.g., *Anopheles gambiae* mosquitoes : Fontaine *et al.* 2015; *Xiphophorus* fishes : Cui *et al.* 2013; African lake cichlids : Meier *et al.* 2017; Caribbean *Cyprinodon* pupfishes: Richards & Martin 2017; *Heliconius* butterflies :Martin *et al.* 2013) that recognized a prominent role of introgressive hybridization as a source of reticulate phylogenies, besides ancestral shared polymorphism.

At some GBS loci, Kerguelen mussels possessed alleles characteristic of both *M. edulis* and *M. galloprovincialis* or *M. trossulus* indicating polymorphism for Northern species-specific alleles in the Kerguelen. Importantly, these loci did not depart from Hardy-Weinberg and linkage equilibrium as shown by an ADMIXTURE analysis (Figure S7) in which the Kerguelen mussels appeared as a well demarcated panmictic cluster. Therefore, contrary to what is known in Northern hybrid zones (Bierne *et al.* 2003), there is no evidence of reproductive barriers impeding admixture in the Kerguelen Islands. Several hypotheses can be proposed: (i) a weaker reproductive barrier between Northern backgrounds at the time of contact in the South; or (ii) an insufficient barrier to gene flow under the demographic and environmental conditions, specifically strong genetic drift, high-potential for hybridization in this small isolated island, or a strong demographic asymmetry between the native and the introduced populations. The first hypothesis relies on the importance of Dobzhansky-Müller incompatibilities for reproductive isolation (Coyne & Orr 2004), and may explain how *M. edulis*, *M. galloprovincialis* and *M. trossulus* alleles at different loci can co-exist into a single Southern population that did not evolve their incompatible interactors, as opposed to the Northern populations. The second hypothesis is based on the fact that the outcome of hybridization can be highly dependent on the demographic context.

Finally, by collecting mussels from contrasted habitats, we demonstrated significant differentiation across 35 sampling sites at the scale of the island, and at a smaller scale between geographically close sites, especially within the Gulf of Morbihan. As found by Gérard *et al.* (2015) at Glu-5′, and previously at allozyme loci (Blot *et al.* 1989), the strongest structure was observed between the North-South coasts and the Gulf of Morbihan, with a genetic break between RdA and PAF at the three most differentiated loci. The fine-scale genetic structure observed in such a high-dispersal species as mussels is at first sight at odds with selective neutrality. So, we explicitly tested the role of habitat heterogeneity in explaining this differentiation. Our RDA analysis shows that genetic variation was associated with habitats, even after controlling for spatial effects; and the most important factors were the presence of *Macrocystis* kelps, substrate type and slope. Despite being low, this significant habitat-driven genetic differentiation could suggest a role of selection. A caveat is that spatial distance between sampling sites may be better described by oceanographic distance than geographic coordinates, and so we may be overestimating the influence of habitats.

Firstly, genetic-environment associations could be due to local adaptation of the mussels opposed by gene flow between habitats. Accordingly, we observed a significant correlation between the presence/absence of *Macrocystis* and the average foreign allele frequency at the four most differentiated loci. This points toward a primary effect of *Macrocystis* presence/absence, which is a keystone species in marine ecosystems that forms kelp forest serving as substrate and refuge for many molluscs species, including *Mytilus* (Adami & Gordillo, 1999), in areas exposed to wave action. Nevertheless, the RdA site, which has the lowest foreign allele frequency, is not occupied by *Macrocystis* kelp, weakening this local adaptation hypothesis.

Alternatively, the consistent genetic patterns observed across several physically unlinked loci indicate the possible existence of two genetic backgrounds maintained at the scale of the island. We propose, and illustrate by simulations, that a genetic background related to *M. edulis* is trapped at RdA and surrounded by another genetic background related to Chilean *M. platensis* mussels and present everywhere else. The enclosed location of the genetic background at RdA explains that it is strongly introgressed at most markers and thus hard to detect. However, the physical barrier to dispersal between sites PAF and RdA produces a clear genetic break on the West side of the contact. Introgression between the two backgrounds generates gradients in allele frequencies, which are better correlated with habitat variation than geographical distance. The foreign allele (as defined by its frequency in the *M. galloprovincialis* Atlantic population of Iberian Coast) tended to be at higher frequency in shallow sites sheltered from the influence of open marine waters with a low salinity and flat-sandy bottoms, mainly in the inner part of the Gulf of Morbihan. These sites are characterized by an absence of *Macrocystis* kelp beds, as opposed to exposed rocky shores. Port aux Français (PAF) is also the harbour where ships arrive and it is the best place for the arrival of non-native genetic backgrounds. Interestingly, *M. galloprovincialis* alleles are found more frequent in exposed, rather than sheltered sites in the hybrid zone between *M. edulis* and *M. galloprovincialis* in Europe, which would suggest inverted genetic-environment associations between hemispheres. Inverted genetic-environment associations are predicted by the coupling hypothesis (Bierne *et al.* 2011), which proposes that genetic-environment associations can easily be revealed by intrinsically maintained genetic backgrounds in linkage disequilibrium with local adaptation genes, and that the phase of the disequilibrium can reverse when contacts are replicated as could have happened in Southern Hemisphere mussels. Overall, these findings reinforce the idea that genetic variation can be maintained at fine geographical scales in high-dispersal organisms, as recently shown in Chilean mussels (Araneda *et al.* 2016) or in passerine birds (Szulkin *et al.* 2016, Perrier *et al.* 2017). In these examples however the link with a possible history of admixture has not been investigated.

Although we had the hypothesis that admixture events could be an explanation of the micro-structure observed in the Kerguelen (Gérard *et al.* 2015), we could not know which backgrounds were in interaction on the sole basis of the GBS data of a single sample. However, our procedure of identifying SNPs that were both polymorphic in the Kerguelen and highly differentiated between Northern Hemisphere species proved to be an interesting way to enrich for loci able to reveal the micro-geographic structure in the Kerguelen. Luckily the sample we used for the GBS analysis (sample "ker-GBS") was localised in the introgression cline, and this can also explain why the enrichment procedure was successful. This is exemplified with the genealogy around locus X10 (Figure S8), which shows that a SNP that differentiates *M. edulis* from other Northern species, and was polymorphic in the Kerguelen Islands, is able to reveal the cline of introgressed heterospecific allele we observed in the island.

In this work, we showed that the most differentiated SNPs in the Kerguelen and those that most strongly drive the genetic-environment associations are primarily ancestry-informative, suggesting that maintenance of genetic differentiation at a small spatial scale, and possibly adaptation to fine-scale environmental variations in the island, may have been facilitated by admixture and introgression of alleles from Northern species. These foreign alleles may have adaptively introgressed the Southern background in the Kerguelen, but the signal is probably erasing because of recombination between adaptive alleles and our neutral markers, and is also probably further blurred by genetic drift. A central question is whether admixture is a simple source of variation on which local selection can effectively act or if the initial linkage disequilibria between foreign alleles in the donor background are required for the successful emergence of micro-geographic adaptation (or speciation in the case of cichlids,Meier *et al.* 2017) and are maintained rather than built-on. In the case of Kerguelen mussels, the evidence we gained here for the maintenance of linkage disequilibrium are limited and indeed rather support extensive recombination. However our markers have likely lost too much signal to answer the question. Maybe local adaptation is operating at loci linked to the candidate SNPs, but most probably these markers simply better reveal a genome-wide signal of habitat constrained connectivity (Gagnaire *et al.* 2015). Overall, our work underlines the opportunity of using non-equilibrium introgression clines to assess genetic connectivity in natural populations and warns against systematically interpreting genetic-environment association as signal of local adaptation.

## Data Accessibility

**Text S1.** Pairwise FST values between Northern species at the GBS SNPs. (A) FST between Med and Nor; (B) FST between Med and Tva; (C) FST between Nor and Tva; Nor: North Sea *M. edulis*; Med: West-Mediterranean *M. galloprovincialis*; Tva: Baltic Sea *M. trossulus*.

**Text S2.** KASPar genotypes of each individual in the Kerguelen Islands (35 sampling sites), plus those of the additional individuals from other Southern Hemisphere populations (6 sampling sites).

**Text S3.** Individual ancestries estimated with ADMIXTURE for the GBS samples. (A) K=2; (B) K=3; (C) K=4; (D) K=5; (E) K=6; (F) K=7; (G) K=8.

**Text S4.** Individual ancestries estimated with ADMIXTURE for the KASPar samples in the Kerguelen (K=2; defining reference populations as *M. edulis* and Chilean mussels).

**Text S5.** Joint site frequency spectrum (in ∂a∂i format) between the Kerguelen mussels and (A) *M. edulis;* (B) *M. galloprovincialis*; (C) *M. trossulus*.

**Text S6.** Definition of the eight models of divergence used in our inferences with ∂a∂i.

**Text S7.** Neighbour-joining trees of the 395 retained GBS sequences.

**Dryad doi: 10.5061/dryad.6k740.** DNA sequences and VCF files including GBS genotypes of each individual in the ten GBS-typed populations (nine Northern populations and eight Kerguelen mussels).

## Supplementary Information

Supplementary Information M&M

Supplementary Information Tables (S1 – S12)

Supplementary Information Figures (S1 – S8)

Supplementary Information Texts (S1 – S7)

## Author Contributions

*Data acquisition*: A. Haguenauer, A. Weber and K. Gérard. *Data analysis*: C. Fraïsse, A. Chenuil and N. Bierne. *Writing*: C. Fraïsse, A. Chenuil and N. Bierne. *Conceptualization*: C. Fraïsse, A. Chenuil and N. Bierne. *Funding acquisition*: A. Chenuil and N. Bierne.

## Acknowledgements

Jean-Pierre Féral and Christian Marschal provided the eight Kerguelen specimens used for GBS during the PROTEKER campaign (Programme IPEV n°1044). Other Kerguelen samples (>600) were collected during scientific program IPEV-MACROBENTHOS n° 195 (1999-2003) by the technical volunteers (VATs) from IPEV missions 49–53 in Kerguelen. This work was funded by the Research Network GDR 3445 cnrs ifremer MarCo and the Agence Nationale de la Recherche (HYSEA project, ANR-12-BSV7-0011). This is article 2015-130 of Institut des Sciences de l’Evolution de Montpellier. This preprint has been reviewed and recommended by Peer Community In Evolutionary Biology (https://dx.doi.org/10.24072/)

